# Remoteness sensitive theta network dynamics during early autobiographical memory access

**DOI:** 10.64898/2025.12.12.694026

**Authors:** Maria Carla Navas, Ignacio Ferrelli, María Eugenia Pedreira, Rodrigo S. Fernández, Luz Bavassi

## Abstract

Autobiographical memory (AM) is a crucial aspect of human cognition that aids in social understanding, decision-making, and future planning. Its inherent diversity provides a valuable framework for exploring the neural dynamics underlying the retrieval of memories of varying ages. This study aimed to determine whether the access phase of AM retrieval contains neural signatures sensitive to the remoteness of these memories. Forty-one participants engaged in an AM retrieval task while their brain activity was recorded using electroencephalography (EEG). We measured spectral power and time-resolved Granger Causality (GC) during the first 1500 ms following the onset of memory recovery. The results showed that remote autobiographical memories led to a significant increase in midline fronto-central theta power (6.5–8 Hz) around 900 ms after retrieval, compared to recent memories. Additionally, GC analyses revealed that both recent and remote memories showed an anterior-to-posterior flow of information, but the remote network present a denser and more efficient connectivity pattern in the theta band between 800 and 1100 milliseconds. These findings suggest that the remoteness of memories influences the early stages of autobiographical memory access. They also highlight theta power and its directed interactions as temporally precise markers of the neural mechanisms that support autobiographical memory retrieval.

## 1 Introduction

Autobiographical memory (AM) is a complex cognitive process that enables individuals to recall personal experiences across their lifespan. In contrast to typical laboratory memory tasks, in which participants encode and retrieve experimentally controlled stimuli, AM paradigms are less constrained due to the inherent variability of individual life experiences. Although this variability poses challenges for experimental control, it provides a unique opportunity to investigate the neural mechanisms underlying the retrieval of memories of different ages [1].

Recalling an AM typically begins with a cue that triggers a controlled search process (the Access period), followed by an Elaboration period during which the memory is reconstructed through mental imagery, as well as the evaluation of its emotional valence and personal relevance [2, 3]. These two phases rely on the coordinated activity of multiple brain regions [4, 5]. Previous studies have suggested a temporal progression of activation, beginning in frontal regions during the initial controlled search and gradually shifting towards posterior temporal and occipital areas as elaboration unfolds [6, 7, 8]. During the Access period, controlled search and effort have been linked to regions such as the medial and right prefrontal cortex (PFC), as well as structures involved in accessing memory traces, including the hippocampus and retrosplenial cortex. As the process progresses into the Elaboration phase, the brain transiently recruits a distributed network, including the anterior medial prefrontal cortex and posterior cingulate cortex, to reconstruct the contextual and subjective richness of the memory [7, 4]. Characterizing the directional interactions within this network is therefore fundamental to understanding how autobiographical memories are accessed and reconstructed.

A fundamental constraint in the study of AM arises from the technical approaches used to investigate it. Most of the evidence supporting the functional progression of activation during AM recall comes from fMRI studies, which provide precise spatial localization but are limited in temporal resolution, typically on the order of seconds. As a result, fMRI is better suited to describe the Elaboration phase, which unfolds over longer timescales. In contrast, this technique limits our ability to characterize the access period, during which memory search is initiated, and fast dynamical interactions between regions may play a crucial role. Consequently, the fine-grained temporal structure and information flow underlying the onset of AM access remain poorly understood.

Brain oscillations play a central role in memory processes [9]. A substantial body of evidence links theta-band activity (4–8 Hz) to memory encoding and retrieval across a wide range of tasks [10, 11, 12, 13, 14]. In particular, increases in theta power have been associated with successful memory retrieval and with preparatory processes preceding a memory response, suggesting a role in cognitive control and decision-making [15]. In the context of autobiographical memory, previous studies have reported distinct oscillatory patterns across frequency bands. For instance, Knyazev et al. showed that AM retrieval is associated with increased alpha (8–12 Hz) and beta (12–30 Hz) power, alongside decreased delta (2–4 Hz) activity, and reported differential modulation of theta power within the Default Mode Network depending on memory vividness [16]. Similarly, Imperatori et al. observed increased delta-band (0.5–4 Hz) power in frontal and medial cortices during AM retrieval [17]. In contrast, Nicolás et al. reported a decrease in theta power around 1000 ms over fronto-central regions when comparing personal versus unfamiliar events [18]. Taken together, these findings highlight the involvement of multiple frequency bands in episodic and autobiographical memory retrieval and suggest that spectral power may serve as neural markers of autobiographical memory retrieval.

Another key aspect of AMs is how their representations evolve and reorganize over time, a process that significantly shapes personal identity and world interpretation. Considering that memory transformation over time comprises a cortical re-organization, the neural mechanisms underlying the retrieval have to be sensitive to the time passed. In this context, AM enables the study of neural correlates of memories of a broad extent of remoteness. Markers of AM age have been reported across different domains: Analysis of speech from free-recall interviews has shown that older AMs tend to contain fewer episodic details and instead rely more heavily on semantic or abstract information [19]. Recent AMs are typically rich in sensory and contextual details, imparting some freshness to our mental representations. In contrast, remote AMs are often reconstructed more schematically, drawing on generalized knowledge or gist-like representations [19, 20]. Also, different patterns of neural representations were found for two-week-old and ten-year-old AM using high-resolution fMRI and multivoxel pattern analysis (MVPA) [21]. The ventromedial prefrontal cortex (vmPFC) showed greater activity in the oldest AMs. At the same time, the hippocampus was involved in both types of memory, with a spatial dissociation indicating that the posterior hippocampus was more strongly associated with the oldest AMs [21]. Besides, Sheldon and Levine, comparing AMs of 1.5 years and of some months, proposed that vividness is a primary factor modulating both the activity and functional connectivity of a brain network involving the hippocampus, exerting a stronger influence than the age of the memory itself. However, subtle effects related to memory age were also observed; older AMs showed activation in the right frontal cortex and left temporal cortex, while recent AMs were associated with the right precuneus and the posterior part of the left hippocampus [22]. A large-scale fMRI meta-analysis by Boccia et al. (2019) provides compelling evidence for this neural reorganization, using classification criteria that distinguish between memories occurring within the last year and those from more distant periods (more than five years ago) [23]. Their findings demonstrate that, while a broad network spanning the occipital to frontal lobes is generally involved in AM, the specific nodes recruited vary with memory remoteness. Memories from the more recent past tend to engage a network including the angular gyrus, dorsomedial prefrontal cortex, and hippocampal-parahippocampal regions. In contrast, retrieval of memories from more distant periods is more strongly associated with activity in the posterior cingulate cortex (PCC), suggesting a shift toward hippocampal-independent storage in cortical hubs. This network-level perspective highlights the importance of examining how information flows across distributed cortical regions, particularly during memory access, to understand how the brain navigates memories of varying remoteness.

Remembering relies not only on localized activity within specific brain regions [24, 8], but also on fast, moment-to-moment interactions across distributed networks. Intracranial EEG (iEEG) studies have shown that both encoding and retrieval involve sub-second fluctuations in functional connectivity, with distinct patterns reinstated during successful episodic recall [25]. These findings suggest that memory access depends on transient shifts in information exchange across brain regions rather than on a purely sequential progression of activation. This perspective highlights the need for analytical tools that capture both the directionality and temporal structure of network interactions. In this sense, Granger Causality (GC), a statistical measure of directed influence between time series [26], provides an appropriate framework for quantifying information flow and characterizing how neural systems coordinate during the early stages of AM access.

Together, this evidence highlights the dynamic and distributed nature of autobiographical memory access and underscores the need for approaches that capture both its temporal structure and directed interactions. In particular, because remote memories are typically less detailed and rely more on reconstructive and schematic processes, their retrieval may require greater large-scale coordination and top-down control. In this context, we aimed to identify neural markers sensitive to memory remoteness during the Access phase by combining exploratory spectral analyses with directed functional connectivity measures. To this end, we developed an experimental protocol designed to capture the fine-grained temporal dynamics of the access period (0–1500 ms). While we explored multiple frequency bands (2–30 Hz), high *θ*-band activity emerged as a key candidate for distinguishing recent and remote memories. Accordingly, we applied frequency-resolved Granger Causality (GC) to characterize the temporal directionality of information flow associated with this neural marker.

## 2 Methods

### 2.1 Participants

41 undergraduate and graduate students, aged from 19 to 33 years old (*mean* = 24.6, *SD* = 3.9; 21 female), participated in the study. All participants self-reported no history of neuropsychiatric conditions. Data from 8 participants were discarded due to noisy neurophysiological recordings. The exclusion criterion was the presence of excessive noise in 10% or more of the channels for prolonged time periods (≈ 20% of the total recording period), as determined by visual inspection.Data are reported for the remaining 33 participants. All participants provided written consent of the Hospital Lanari, in accordance with the Declaration of Helsinki.

### 2.2 Stimuli

The stimuli were designed to elicit neutral autobiographical memory (AM) events, specifically targeting the recall memories of varying remoteness. To enhance the distinction between memory age within our experimental population, we adapted the stimulus set initially developed by [27].

### 2.3 Autobiographical Memory protocol

Each participant completed 24 trials, 12 of which corresponded to *“the first time”* AM events and 12 *“the last time”* AM events. These trials were randomly selected from a dataset of neutral events and presented in a randomized order (*N* = 37; stimuli translated to English are in the supplementary material). Note that the original stimuli are in Rioplatense Spanish. In each trial, participants were cued to retrieve a specific memory via an on-screen sentence such as *“Remember the FIRST TIME you cooked by yourself”* or *“Remember the LAST TIME you took a bus”*, targeting later and earlier memories, respectively. The Access Period is the time spent attempting to access that memory. This Access period was self-paced, meaning that every participant spent as much time as required to access the cued memory. Once the memory came to mind, participants pressed the space bar and mentally elaborated on the event for 8 seconds; this phase is referred to as the Elaboration period ([7, 3, 28, 27, 29]). Afterward, participants provided subjective ratings of the memory, including its emotional valence from very negative to very positive (-2 to 2), level of detail from no detail to a lot of detail (1 to 5), access difficulty from no difficulty to a lot of difficulty (1 to 5), perceived importance from no importance to a lot of importance (1 to 5), and estimated time of occurrence from today to more than 3 years ago using a 7-point Likert scale: 1 (today), 2 (one week), 3 (one month), 4 (Three months), 5 (1 year), 6 (3 years) and 7 (more than 3 years). Participants could select any intermediate value on the continuum of memory-remoteness. If participants were unable to recall a memory, they pressed the ‘s’ key to skip the trial. Figure 1 provides a schematic representation of the experimental protocol. While vividness is a commonly used measure of memory richness in AM studies, there is no single Spanish term for it. We decomposed this construct into multiple subjective ratings to provide a more nuanced and comprehensive understanding of the characteristics of each AM.

**Figure 1.**
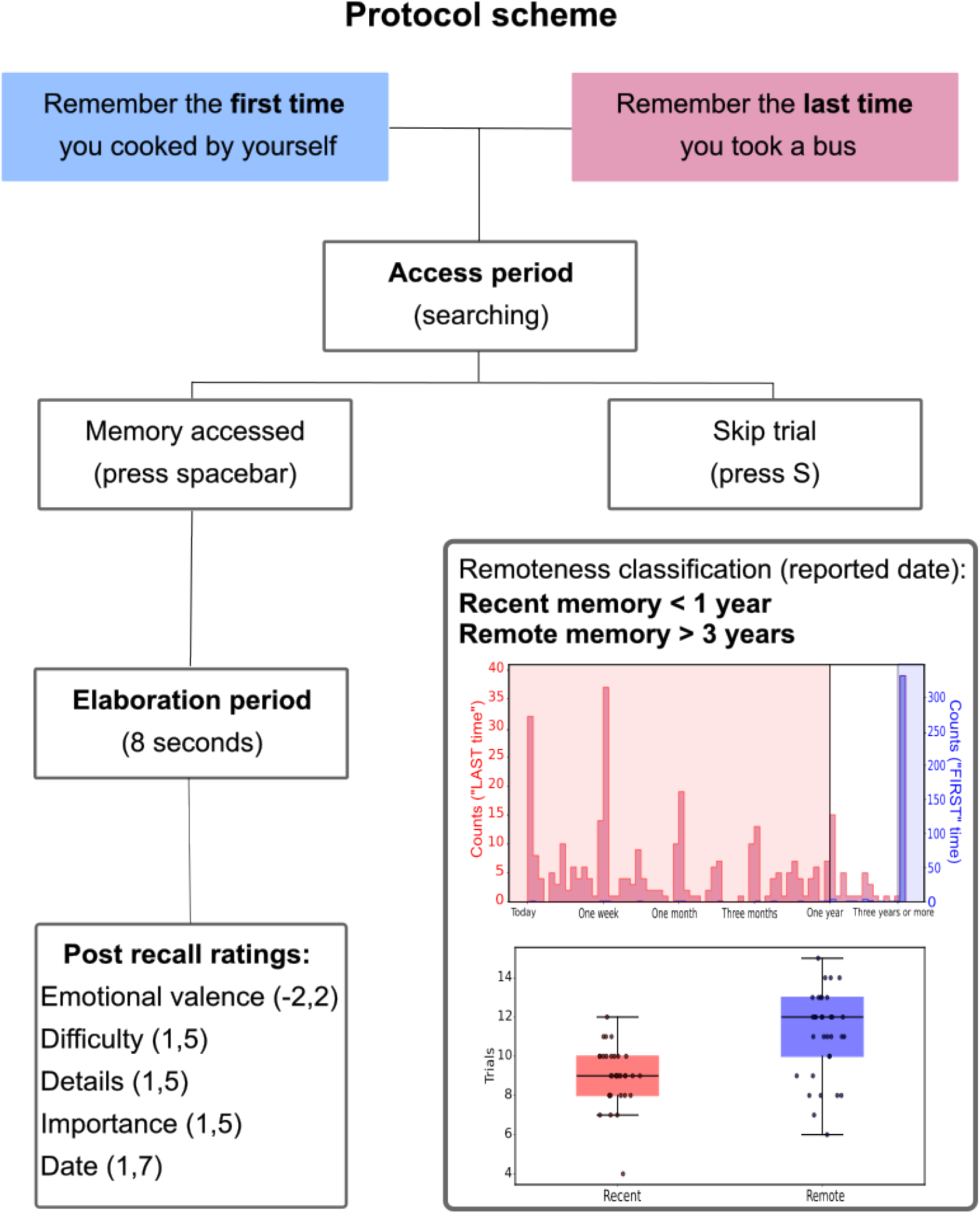
Overview of the Autobiographical Memory task. Participants were cued to recall either a remote AM (blue) or a recent AM (red). The appearance of the cue on screen marked the beginning of the Access period, which lasted until the participants either pressed the space bar once the target memory was accessed (left) or skipped the trial (right). Right after pressing the spacebar, the Elaboration period started: Participants were asked to mentally elaborate the memory with as many details as possible during 8 seconds. After the elaboration, they had to rate that memory in the following categories: level of detail recalled, level of difficulty in accessing the memory, emotional valence of the memory, importance of the event, and how long ago the event occurred. Posteriorly, the reported date was used to classify the memories. The histogram illustrates the distribution of self-reported memory age based on a 7-point Likert scale. Participants used a continuous slider anchored at 1 (today), 2 (1 week), 3 (1 month), 4 (3 months), 5 (1 year), 6 (3 years), and 7 (more than 3 years). The pink histogram represents the “FIRST” time stimuli, while the light blue represents the “LAST” time ones. The red and blue shaded regions indicate the boundaries for categorical assignment: memories rated ≤ 5 (up to 1 year) were labeled as *Recent*, while those rated ≥ 6 were labeled as *Remote*. Anchors 6 and 7 were collapsed into a single one. To maximize temporal contrast, trials in the intermediate 1–3 year interval (5 *<* value *<* 6; *N* = 54) were excluded. Boxplot of the number of trials per condition per participant. In total, 664 trials were included in the analysis (*N*_*rem*_ = 367, *N*_*rec*_ = 297), with an average of 11 ± 2.2 remote and 9 ± 1.5 recent trials per participant.

After data collection, we assigned Recent and Remote labels based on participants’ self-reported timing. As ratings were provided using a continuous slider bar, which allowed participants to select intermediate points between the integer anchors, the resulting scale was non-homogeneous, as the temporal intervals represented between anchors were not uniform (see Figure 1). To ensure clear temporal contrast and minimize ambiguity, events rated up to 1 year (values ≤ 5) were labeled as Recent. In contrast, those occurring 3 years or more prior (values ≥ 6) were labeled as Remote. Trials falling within the intermediate 1 to 3 years interval (values between 5 and 6, total *N* = 54 trials) were excluded from the analysis to maintain a distinct separation between categories. This yielded a total of 664 trials (*N*_*rem*_ = 367 and *N*_*rec*_ = 297). On average, each participant summed 11 ± 2.2 remote trials (range: 6–15) and 9 ± 1.5 recent trials (range: 4–12) after data pre-processing and cleaning.

### 2.4 Equipment

Participants sat comfortably in a chair while stimuli were presented on a screen positioned 60 cm in front of them. The experimental paradigm was developed and presented using custom software written in PsychoPy v2023.2.3 [30]. Brain activity was continuously recorded throughout the task using an Akonic EEG device (https://www.akonic-argentina.com/), with a sampling rate of 256 Hz and 30 EEG channels arranged according to the international 10-20 system. Two additional reference electrodes were placed bilaterally on the mastoids. Behavioral responses were collected via keyboard input. Each response triggered a synchronization marker sent to an external EEG channel via an Arduino Nano (http://www.arduino.cc/), an open source electronics prototyping platform, enabling a temporal resolution of 1 ms.

### 2.5 Data Analysis

#### 2.5.1 Behavioral data

Behavioral data were analyzed using custom software implemented in Python and R. Access times for Remote and Recent AMs ranged from 1900 ms to 65000 ms and were compared using a nonparametric Mann-Whitney U test. We also statistically compared differences in emotional valence, level of detail, access difficulty, and perceived importance between Remote and Recent AMs using Wilcoxon rank-sum tests. Then, we explored the relation between these measures. To this end, we computed the Spearman correlation among them.

To identify the most relevant behavioral predictors of AM age (Recent vs Remote), we implemented a regularized Logistic Regression model. Logistic regression is a classification algorithm that models the probability of a binary outcome (“Recent” (*Y* = 1) or “Remote” (*Y* = 0)) as a function of a set of predictors *X* (detail, importance, difficulty and emotional valence):

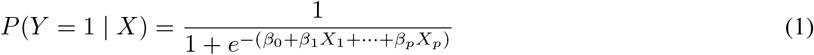

As our behavioral predictors presented multi/collinearity, we incorporated the Elastic Net regularization, which combines *L*_1_ (Lasso) and *L*_2_ (Ridge) norms to perform shrinkage and variable selection simultaneously [31]. It does so by minimizing a cost function (*L*, the negative log-likelihood for the binomial family) where parameter *λ* controls the strength of the regularization and *α* the balance between *L*_1_ and *L*_2_ penalties. To this end, we conducted a grid search using the train function of the caret package in R [32] to optimize the model’s hyperparameters. We explored 100 log-scaled values of the regularization parameter *λ* (ranging from -4 to 0) and 11 values of the mixing parameter *α* (ranging from 0 to 1) using the glmnet method [33]. We evaluated the model performance through 5-fold stratified cross-validation, and the optimal *α*-*λ* combination was selected based on the area under the ROC curve (AUC). The best-performing Elastic Net model was then validated on a held-out test set, with classification performance assessed via overall accuracy and confusion matrix. To further evaluate the stability of the selected model coefficients, we performed nonparametric bootstrapping (1000 iterations) on the training set. This procedure involves refitting the selected model on each of the 1000 resamples to estimate the distribution and variability of the coefficients. Predictors were identified as statistically reliable if their 99% bias-corrected and accelerated (BCa) confidence intervals did not include zero. This criterion ensures that the relationship between the predictor and the outcome remains consistent in direction, with a 99% probability that the true coefficient is non-zero. The BCa was estimated using the boot package in R. Finally, our analysis concluded that *α* = 1 and *λ* = 0.009.

#### 2.5.2 EEG data preprocessing

EEG data were band-pass filtered between 1 and 35 Hz using a zero-phase 8th-order Butterworth IIR filter to eliminate slow oscillatory trends and high-frequency noise. Then, the data were re-referenced to the average activity using MNE’s projection method. Noisy channels were identified through visual inspection and subsequently interpolated. In total, 22 subjects had interpolated channels, with a maximum of 3 channels interpolated per subject (10%). We applied Independent Component Analysis (ICA) to look for eyeblink artifacts, we excluded 2 ICA compenents on average per participant. After a visual inspection, we removed the components that reflected these physiological responses. ICA was computed using the FastICA algorithm [34], as implemented in the MNE-Python toolbox (https://mne.tools/stable/index.html) [35].

We epoched the raw data into two 2000ms sets of epochs, aligned to the presentation of the “Remember…” screen (stimulus-locked epochs) and to the keypress when memory was accessed (centered on the response). The epochs aligned to the stimulus began 500 ms before screen onset and extended until 1500 ms after its appearance. Although the minimum extent of the Access period was 1900 ms, we focused the EEG analysis on the first 1500 ms to reduce potential contamination from motor preparation and early elaboration processes. This precaution is particularly relevant given that spectral and Granger Causality analyses can introduce temporal smearing, causing later activity to influence earlier time windows. Complementary, the epochs aligned with the response began 1500ms before the key was pressed and extended by 500ms (response-locked epochs).

Then, we performed trial rejection using the Autoreject algorithm [36], which automatically estimates an optimal global amplitude threshold for rejecting noisy epochs. This threshold is determined via a data-driven cross-validation approach. For a range of candidate thresholds, the algorithm splits the data into k folds. It compares the median signal from the training set (after rejecting bad trials at each candidate threshold) with the median signal from the test set. The error between these signals is quantified using the Frobenius norm, and the threshold that minimizes this error is selected. After trial rejection, the total number of trials was 664 trials (*N*_*rem*_ = 367 and *N*_*rec*_ = 297). On average, each participant summed 11 ± 2.2 remote trials (range: 6–15) and 9 ± 1.5 recent trials (range: 4–12).

#### 2.5.3 Time-frequency analysis

For spectral analysis, a Morlet wavelet transform was applied to each epoch using the FieldTrip toolbox in Matlab https://www.fieldtriptoolbox.org/ [37]. Spectral power was estimated between 2 and 30 Hz, with a frequency resolution of 0.5 Hz, using a 5-cycle wavelet at each frequency. Time-frequency decomposition was performed on the full epochs (stimulus/response locked-epochs) of the Access period. Spectral power was normalized using the average power across time for each participant, frequency and channel. Finally, the resulting power values were converted to decibels (dB).

#### 2.5.4 Cluster-based time-frequency comparison

To compare spectral power between the Remote and Recent Access periods, we performed a nonparametric cluster-based permutation test using the FieldTrip toolbox in MATLAB. We employed a within-subjects design and used a dependent-samples t-test to investigate differences between the two conditions. Cluster correction for multiple comparisons was performed using the Monte Carlo method with 1500 permutations. The forming cluster’s threshold was set at p < 0.05, with a minimum of two neighboring channels. A two-tailed test was conducted, and the statistic for cluster-level inference was the sum of t-values [37]. We performed this statistical analysis on all EEG channels and across the full epoch (stimulus and response - locked epochs) for the frequency bands *δ* (2-4), *θ* (4-8), *α* (8-12 Hz), *β*_1_ (12-20 Hz), *β*_2_ (20-30 Hz).

#### 2.5.5 Directed connectivity analysis

##### EEG processing

To estimate directed functional connectivity in the frequency domain, we first processed the EEG signals before applying Granger Causality (GC) analysis (Figure 2). To ensure the stationarity required for Multivariate Autoregressive Model (MVAR) modeling, we performed a two-step processing procedure. First, the ensemble mean (ERP) was calculated for each subject and separately for each experimental condition. This condition-specific ERP was then subtracted from each individual trial using a point-by-point approach (bsxfun in MATLAB) across the entire epoch. This procedure removes deterministic trends that could otherwise lead to spurious connectivity estimates. Then, we segmented the EEG signals into 400 ms sliding time windows with 10 ms steps, from -500 to 1500 ms relative to the screen presentation. Each 400ms segment was detrended and standardized using a z-score to improve stationarity. Next, we concatenated those segments for all Recent or Remote stimulus-locked epochs of each subject.

**Figure 2:**
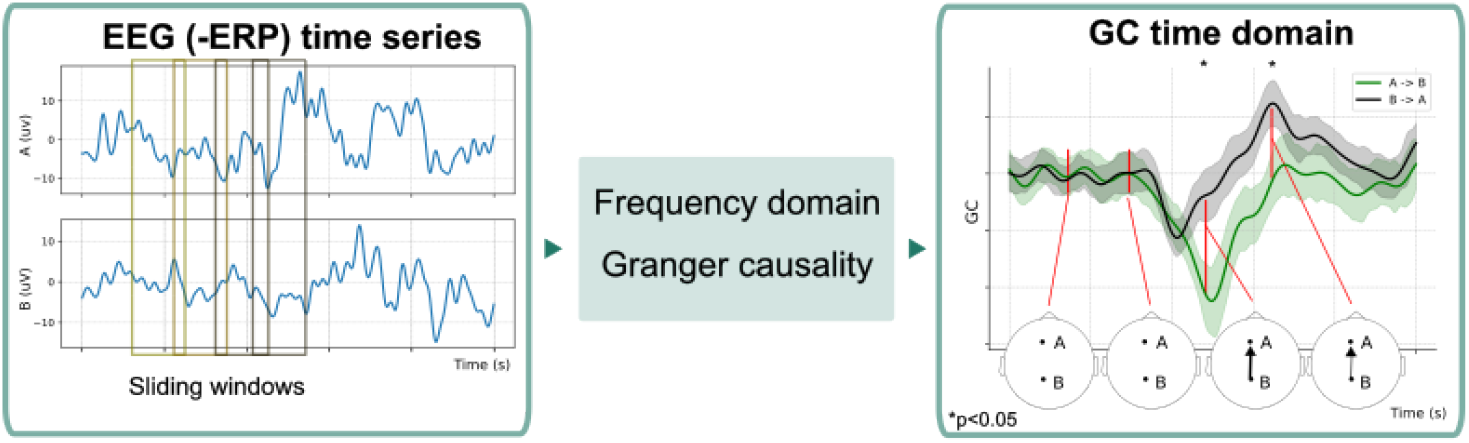
Schematic representation of the building of high-*θ* GC networks. First, we subtracted the ERP from each channel time series. The data were then segmented into 400 ms sliding windows with 10 ms steps. Each segment was subsequently z-scored and detrended prior to computing frequency-domain Granger Causality (GC). The resulting time series represent GC values for the directed influences from channel A to B (green) and from B to A (black) in the high-*θ* band. For each time window, we performed a Wilcoxon signed-rank test across subjects on the GC values for each channel pair to assess the dominant direction of information flow. Shaded areas represent the standard error (SE). When *p <* 0.05, the edge weight was defined by the corresponding GC value.

##### Frequency-domain Granger Causality

To estimate directional statistical relation between electrodes in the frequency domain, we used frequency resolved GC [38, 39], as implemented in the BSMART toolbox freely available for download online under the GNU general public license https://brain-smart.org/) [40]. The method is based on fitting bivariate autoregressive models (AMAR) to pairs of EEG channels (A(t),B(t)). Let *X*(*t*) = [*A*(*t*), *B*(*t*)]^⊤^ represent a bivariate stationary process within a time window, where ⊤ denotes the transpose.

The signals are modeled using a *p*-th order autoregressive process:

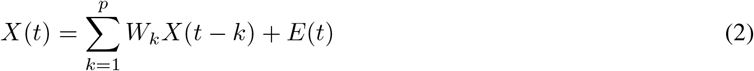

where *W*_*k*_ ∈ ℝ^2×2^ are the coefficient matrices and *E*(*t*) is a zero-mean white noise process with covariance matrix Σ.

The model parameters are estimated using the Levinson-Wiggins-Robinson (LWR) algorithm [41] via the Yule Walker equations.

Once the model is estimated, it is projected into the frequency domain via the transfer function:

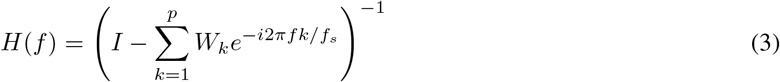

where*f*_*s*_ is the sampling frequency.

The spectral density matrix *S*(*f*) is then computed as:

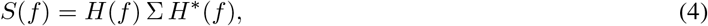

where *H*^∗^(*f*) denotes the conjugate transpose of *H*(*f*) and Σ is the noise covariance matrix of the bivariate autoregressive process. This matrix represents the covariance of the white noise innovations *E*(*t*), that is, the residuals or errors that the model cannot predict from past values.

The frequency resolved GC from channel *j* to channel *i* is defined as:

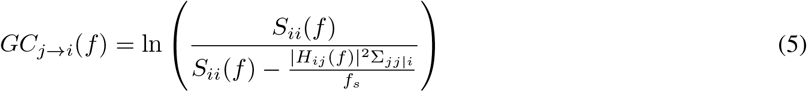

where *S*_*ii*_(*f*) is the power spectral density of channel *i, H*_*ij*_(*f*) is the transfer function component from *j* to *i*, Σ_*jj* |*i*_ is the residual noise variance when excluding the influence of channel *j*.

A value of *GC*_*j*→*i*_(*f*) *>* 0 indicates that past activity in channel *j* 4 helps predict the activity in channel *i* at frequency *f*, a *directed Granger-causal influence* from *j* to *i*, while *GC*_*j*→*i*_(*f*) ≈ 0 suggests no predictive information from *j* to *i* at that frequency.

The model order (*p*) was determined by evaluating the Bayesian Information Criterion (BIC) across a range of potential orders (4 to 117ms). Our results showed that BIC values decreased sharply, reaching a plateau between 80 and 98ms. This choice provides a model that captures the signal’s directional dynamics while avoiding overfitting. Furthermore, maintaining a relatively short model order prevents excessive parameter inflation that would occur if the order were extended to cover full cycles of the low frequencies, thereby preserving the temporal resolution required for analyzing early memory access stages. In conclusion, the model order was set to 80 ms (*p* = 20).

##### Functional connectivity: Directed Weighted Networks

After fitting the autoregressive model, we obtained GC time series for each pair of EEG channels identified in the cluster-based analysis (Fp1, Fp2, Fpz, Fz, Cz, Pz, FC1, FC2, CP1, CP2, AFz; Figure 4), separately for each condition (Remote, Recent), frequency, and participant. We then averaged GC values across the 6.5–8 Hz range to obtain time series in the high-*θ* band, corresponding to the frequency range identified as significant in the previous analysis.

To build the global time-resolved networks (for all participants), we performed Wilcoxon signed-rank tests for each channel pair, time window, and condition to identify moments of statistically significant GC directionality in the high-*θ* across all subjects. Significance was defined at p < .05 (uncorrected), and the direction of the effect was determined by the sign of the test statistic (Z value). Figure 2 is a schematic representation of the building of the GC high-*θ* networks.

Then, we examined the temporal evolution of time-resolved networks by characterizing network density over the first 1500 ms of the access period. Network density was defined as the number of observed links divided by the total number of possible links (*E*_*tot*_ = 72) [42]. To assess differences between conditions, we performed a nonparametric paired Wilcoxon signed-rank test on network density values, treating time points within the access period as repeated observations for each participant.

Subsequent analyses focused on the 800–1100 ms interval, corresponding to the time window in which significant differences between Recent and Remote AM access were observed. For each condition, we aggregated the time-resolved networks within this window. To characterize these networks, we computed the following topological measures [42]:

- **Number of connected nodes (***N* **) and links (***E*).
- **Mean Degree (**⟨*k*⟩): The average number of connections per node:

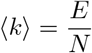
- **Mean Strength (**⟨*s*⟩): The average sum of connection weights (*w*_*ij*_) per node:

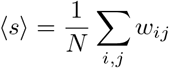
- **Clustering Coefficient (***C*): A measure of local segregation. For directed and weighted networks, we computed the nodal clustering (*C*_*i*_):

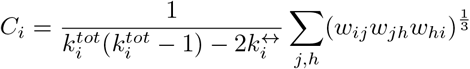

The global clustering for the whole network is the average of all these nodal coefficients:

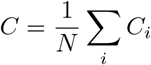

- **Global Efficiency (***Eff* ): A measure of network integration based on the shortest directed path length (*d*_*ij*_) between nodes *i* and *j*:

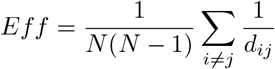

To evaluate whether directed information flow exhibited a spatial bias between anterior and posterior regions, we implemented a subject-specific normalization procedure based on a null model. EEG electrodes were grouped into an anterior cluster (*N* = 7; Fp1, Fp2, Fpz, Fz, AFz, FC1, FC2) and a posterior cluster (*N* = 4; Cz, CP1, CP2, Pz). For each participant, within the 800–1100 ms window, we computed the observed directional flow strength (*obs*) by summing GC values across four categories: anterior-to-anterior, posterior-to-posterior, anterior-to-posterior (*A* → *P*), and posterior-to-anterior (*P* → *A*). To account for potential topological biases arising from unequal electrode counts per cluster, we generated a null distribution for each subject using a permutation procedure (*n* = 1500). In each iteration, electrode labels were randomly reassigned across nodes, and connectivity sums were recomputed for each category. Observed values were then normalized by the mean of the corresponding null distribution

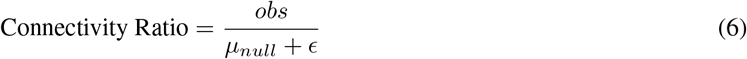

where *µ*_*null*_ denotes the mean of the null distribution and *ϵ* is a small constant to prevent division by zero. A ratio greater than 1 indicates that the observed flow exceeds chance expectations given the network topology.

### 3 Results

#### 3.1 Behavioural: Characterization of recall

This work aims to characterize differences in access to autobiographical memories (AMs) based on remoteness. First, we compared the behavioral auto-reported measures for Recent (less than 1 year) and Remote (more than 3 years) AMs. We explored the duration of the Access period. The access time was defined as the latency from the onset of the cue until participants pressed the space bar to indicate that a specific AM was located. No statistically significant difference was found between Recent and Remote Access periods (Mann-Whitney Z = 1.34, p = 0.18); the median latency of Access period was 7.1 seconds, IQR = 5.4 seconds for Recent AMs and 6.6 seconds, IQR = 4.8 seconds for Remote AMs Figure 3 A.

**Figure 3.**
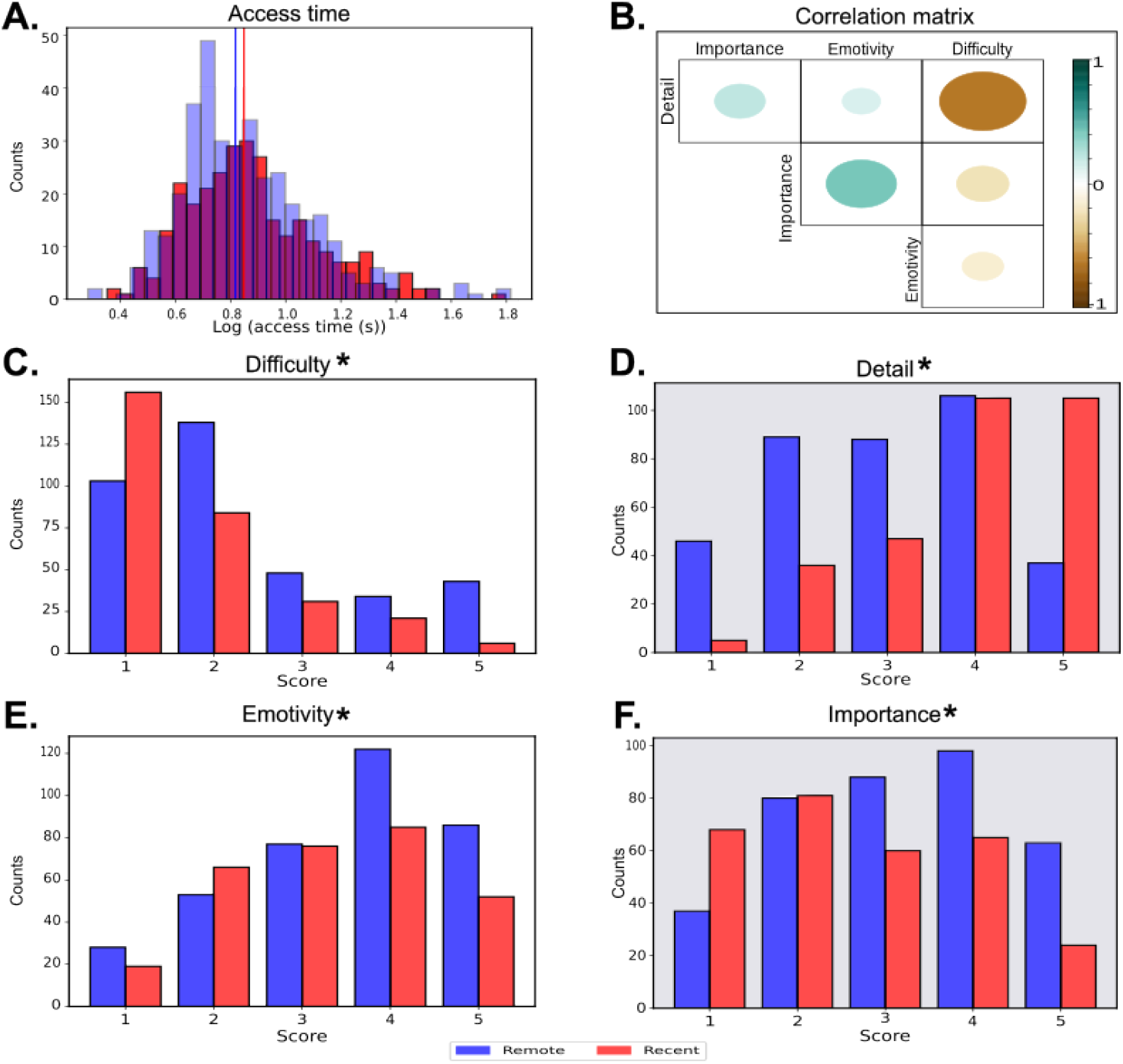
Behavioral results. **A**. Access time histogram for Recent (red) and Remote (blue) trials. Mann-Whitney U = 57791.5, p = 0.18; Vertical lines represent the median latency of access Period for Recent (7.1 seconds) and Remote (6.6 seconds) AMs. **B**. Correlation matrix of the self-reported measures. The colored circles represent the correlation coefficient. **C**,**D**,**E**,**F:** Distribution of responses for the self-reported measures. The gray frame highlights the two measures that are most important for distinguishing between Recent and Remote events, as determined by logistic regression.

Then, we explored the ratings for each self-reported measure: emotional valence, importance of the event, details recalled, and difficulty remembering. We show the distributions of responses for each condition Figure 3 C-F. To evaluate differences in the subjective phenomenological characteristics between Recent and Remote memories, a series of Wilcoxon rank-sum tests were conducted. The analyses revealed significant differences between the two conditions across all evaluated dimensions. Participants reported significant differences in difficulty (*Z* = − 6.57, *p <* .001), emotionality (*Z* = − 2.58, *p* = 0.009), importance (*Z* = − 5.34, *p <* .001) and details (*Z* = 9.51, *p <* .001). Overall, recent memories were rated as less difficult, less emotional, less important, and more detailed compared to remote memories. We examined the associations between behavioral variables to detect linear relations between them. Spearman correlations showed a robust negative association between detail and difficulty (*ρ* = − 0.62, *p <* .001), indicating that memories retrieved with higher levels of detail were also perceived as easier to access. A moderate positive correlation was observed between importance and emotivity (*ρ* = 0.41, *p <* .001), suggesting that participants considered memories with a higher emotional charge to be more significant. Furthermore, significant but weaker correlations were found between importance and difficulty (*ρ* = −0.23, *p <* .001), detail and importance (*ρ* = 0.21, *p <* .001), emotivity and difficulty (*ρ* = −0.14, *p <* .001), and detail and emotivity (*ρ* = 0.12, *p* = .002). (Figure 3 B).

Finally, we performed a classification model to predict AM remoteness as a function of the behavioral self-reported measures. To discard redundant information due to variable co-linearity, we performed an Elastic Net logistic regression:

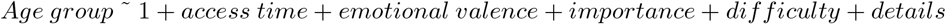

We found that the most important features to differentiate Recent and Remote trials were importance (*β* = -0.56) and details (*β* = 0.75). All the coefficients of the model are shown in Table 1. This shows that despite Remote stimuli being rated as more important, they tended to be recalled with fewer details than Recent ones.

**Table 1:**
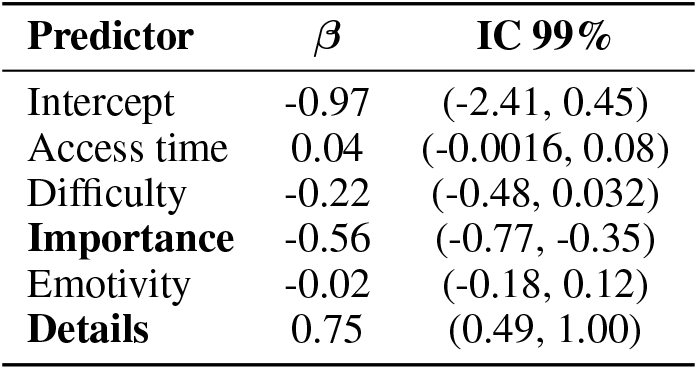
Logistic Regression.

| Predictor | $\beta$ | IC 99% |
| --- | --- | --- |
| Intercept | -0.97 | (-2.41, 0.45) |
| Access time | 0.04 | (-0.0016, 0.08) |
| Difficulty | -0.22 | (-0.48, 0.032) |
| **Importance** | -0.56 | (-0.77, -0.35) |
| Emotivity | -0.02 | (-0.18, 0.12) |
| **Details** | 0.75 | (0.49, 1.00) |

#### 3.2 Neural correlates

First, we visualized the EEG activity during the access period. Figure 4A,B show the voltage time series across all electrodes for stimulus-locked and response-locked epochs, respectively. As expected, the onset of the screen change evoked a rapid response (*<* 200 ms) across all channels in both types of epochs. In the stimulus-locked condition, high-amplitude activity extended over the first 500 ms, likely reflecting visual and reading processes. In contrast, response-locked epochs exhibited a marked decrease in frontal channels approximately 100 ms prior to the keypress, consistent with premotor processes.

**Figure 4.**
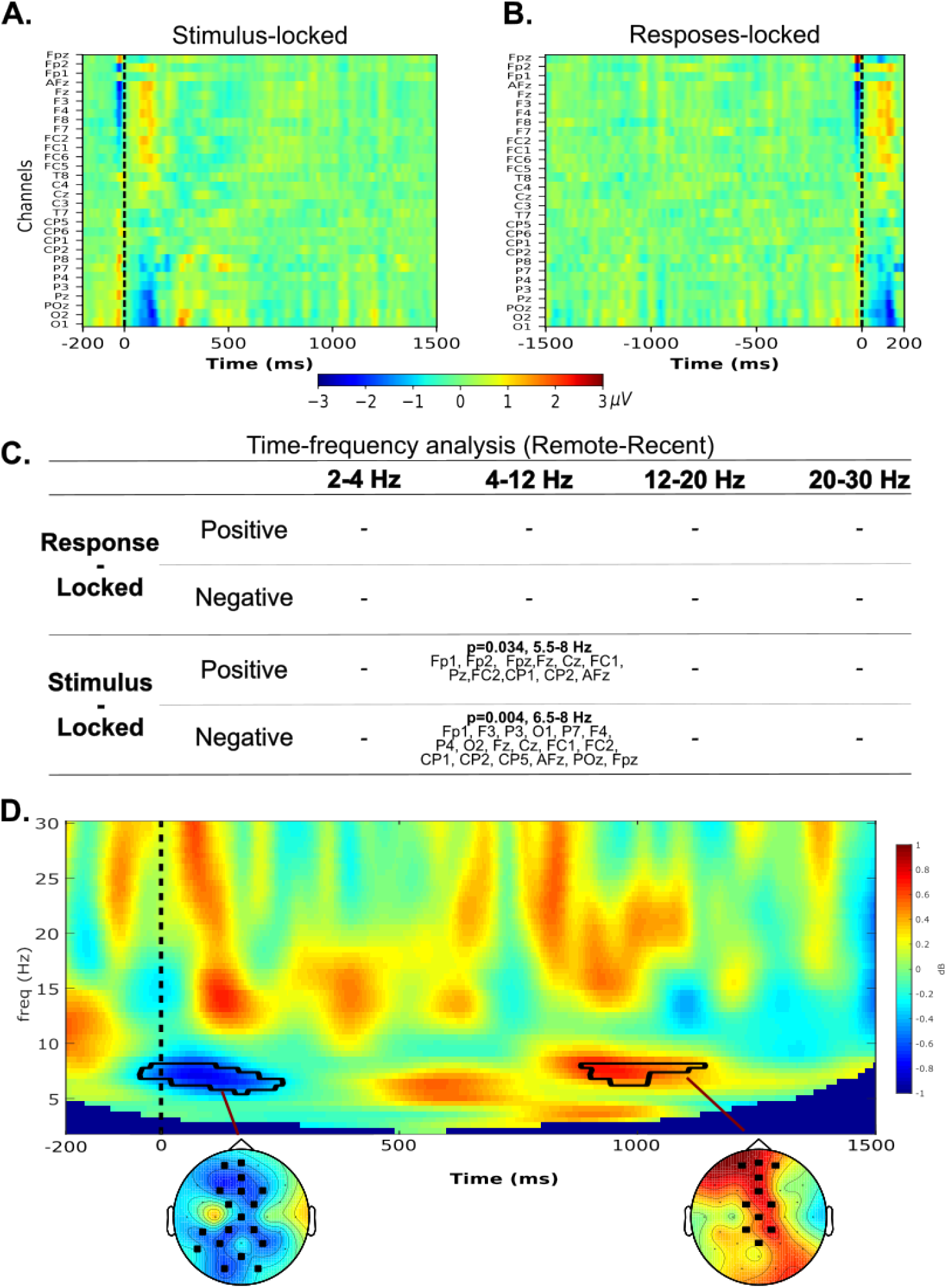
Temporal and spectral dynamics of AM access. **A and B**. Voltage scalp activity for stimulus-aligned and response-aligned epochs. **C**. Table of results of the time frequency analysis per frequency band and stimulus/response-aligned epochs. **D**. Time frequency power for the mean of the channels forming both clusters along the whole stimulus-aligned epochs (cold colors depict Recent>Remote, warm colors represent Remote>Recent). The black trace highlights the significant clusters, the positive cluster. Topomaps displaying the significant clusters found in *θ* band using a non-parametric cluster-based permutation test: A negative one p = 0.004 between 5.5 - 8 Hz from -40 to 250 ms involving channels Fp1’, ‘F3’, ‘P3’, ‘O1’, ‘P7’, ‘F4’, ‘P4’, ‘O2’, ‘Fz’, ‘Cz’, ‘FC1’, ‘FC2’, ‘CP1’, ‘CP2’, ‘CP5’, ‘AFz’, ‘POz’, ‘Fpz’, (left) and a positive one p = 0.038 between 6.5 - 8 Hz from 880 and 1140 ms involving electrodes Fp1, Fp2, AFz, Fpz, Fz, Pz, Cz, FC1, FC2, CP1 and CP2 (right).

##### 3.2.1 Time frequency analysis

Based on previous literature, we examined spectral power to identify neural markers of AM remoteness during the access period. We conducted parallel time-frequency analyses using both stimulus- and response-locked epochs. Differences in power across frequency bands (*δ*: 2–4 Hz, *θ*: 4–8 Hz, *α*: 8–12 Hz, *β*_1_: 12–20 Hz, *β*_2_: 20–30 Hz) were assessed using a cluster-based permutation paired-samples *t*-test (Remote vs. Recent).

Significant differences between conditions were observed only in the stimulus-locked data. Accordingly, subsequent analyses focused on stimulus-locked epochs, as no significant clusters were identified in the response-locked data (Figure 4C). Within stimulus-locked epochs, two significant clusters were identified in the *θ* band (Figure 4 D). The first was a negative cluster (5.5–8 Hz) spanning from -40 to 250 ms and involving a widespread set of electrodes (Fp1, F3, P3, O1, P7, F4, P4, O2, Fz, Cz, FC1, FC2, CP1, CP2, CP5, AFz, POz, Fpz). This time window overlaps with early evoked visual responses (*<* 200 ms), suggesting that the observed differences may reflect low-level perceptual processing rather than memory-related activity. Based on this interpretation, this cluster was not considered a marker of AM remoteness. In contrast, a significant positive cluster was observed in the high *θ* (6.5 and 8 Hz), spanning 880 to 1140 ms, and involving electrodes Fp1, Fp2, AFz, Fpz, Fz, Cz, Pz, FC1, FC2, CP1, and CP2 (Figure 4 D). This cluster reflected greater *θ* power over midline frontal and central regions for Remote compared to Recent AMs during the early stages of memory access.

To evaluate the subjective factors modulating the identified *θ* marker (defined as the average power across the selected channels within the *θ* frequency band during the specific time window of interest), we performed a generalized linear mixed-effects model (GLME). The model included difficulty, emotionality, importance, and level of detail as fixed effects. To account for the non-independence of the data, random intercepts for participants and stimulus numbers were also included.

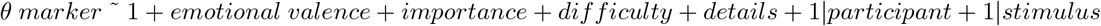

The results shown in Table 2 revealed a significant main effect of question difficulty on *θ* power (p = 0.04), indicating that a higher level of difficulty was associated with increased *θ*. This suggests that *θ* activity in this paradigm is tied to the perceived difficulty of memory access rather than to the details or subjective qualities of the memory.

**Table 2:** Generalized Linear Mixed-Effects Model.

| Predictor | $\beta$ | IC 95% |
| --- | --- | --- |
| Intercept | -3.19 | (-4.18, -2.20) |
| **Difficulty** | 0.17 | (0.01, 0.34) |
| Importance | 0.07 | (-0.07, 0.20) |
| Emotivity | -0.04 | (-0.18, 0.10) |
| Details | 0.11 | (-0.06, 0.27) |

##### 3.2.2 Functional connectivity

To investigate the neural flow of information underlying AM access, we constructed directed connectivity networks using GC in the high-*θ* band for the subset of fronto-central electrodes identified in the previous analysis. We first examined the temporal evolution of network density (Figure 5A). Although the number of links appeared similar across conditions, the Remote AM network exhibited a significantly higher density compared to the Recent AM network (paired Wilcoxon signed-rank test: *Z* = −2.06, *p* = 0.03; Figure 5B).

**Figure 5.**
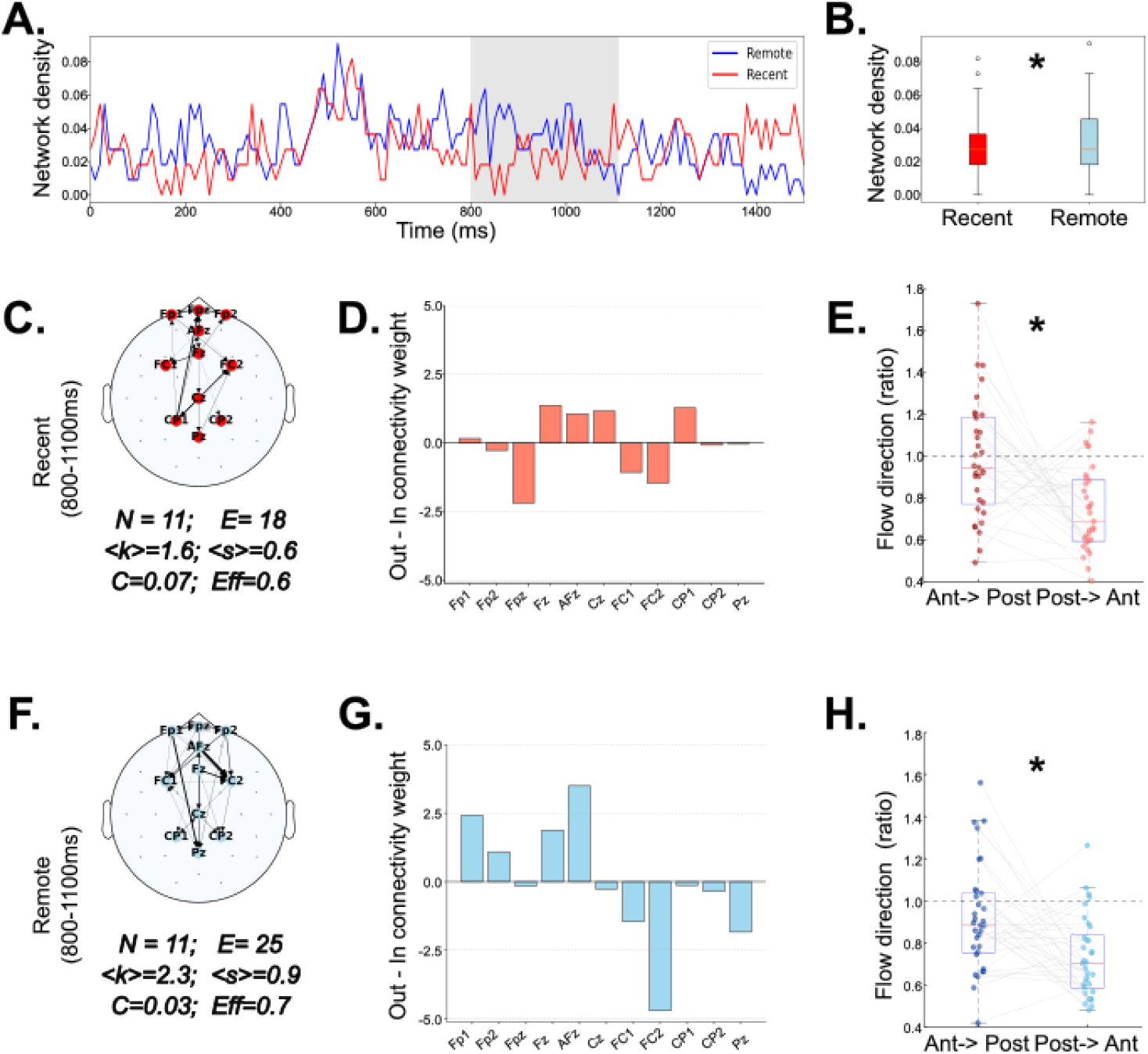
GC *θ* networks. A. Time series of network densities. The highlighted window (800–1100 ms) indicates the period where the difference between conditions reaches its maximum, based on the 95th percentile distribution. **B**. Density per condition over time. Density of the Remote networks is statistically denser than the recent one.(*Z* = − 2.06, *p* = 0.03). **C**. and **F**. Topology of the connectivity networks in the highlighted time window. **D**. and **G**. Difference between out degree - in degree for each channel. **E**. and **H**. GC values for all participants divided into anterior to posterior connections and posterios to anterior. Anterior electrodes: Fp1, Fp2, Fpz, Fz, AFz, FC1, FC2; Posterior electrodes: Cz, CP1, CP2, Pz.

We then focused on the 800–1100 ms window identified in the time–frequency analysis. In this interval, both networks included all nodes, but differed in their connectivity patterns. The Recent network (Figure 5C) comprised 19 connections (E =19), whereas the Remote network (Figure 5F) showed a higher number of connections (*E* = 25), along with increased mean degree (⟨*k*⟩) and mean strength (⟨*s*⟩). The Recent network exhibited slightly higher clustering (*C* = 0.07), suggesting more locally segregated connectivity, whereas the Remote network showed higher global efficiency (*E*_*global*_ = 0.7), indicating shorter paths for information transfer.

Examination of directed connectivity patterns revealed distinct flow organizations. In the Recent network, the main outgoing hubs were located in midline fronto-central regions (Fz, Cz, AFz), with primary targets in bilateral frontal areas (Fpz, Fp1, Fp2) (Figure 5D). In contrast, the Remote network showed a predominant anterior-to-posterior flow, with information propagating from bilateral frontal regions toward parietal midline areas (Figure 5G).

To further assess this directional bias, we analyzed GC values at the participant level within the 800–1100 ms window (Figure 5E,H). In both conditions, the dominant direction of information flow was anterior-to-posterior (Remote: *Z* = 2.47, *p* = 0.013; Recent: *Z* = 2.96, *p* = 0.003), with no significant differences between conditions in the magnitude of this directional effect.

Overall, these results indicate that Remote AMs are associated with a more densely connected and globally efficient high-*θ* network during the access period. While both conditions exhibit a common anterior-to-posterior flow of information, Remote memories engage a more integrated network architecture.

## 4 Discusion

In the present study, we characterized the neural dynamics underlying autobiographical memory (AM) access. Using an exploratory spectral analysis across multiple frequency bands, we identified high *θ*-band activity (6.5–8 Hz) as a sensitive neural marker of memory remoteness. Specifically, Remote AMs elicited greater *θ* power over midline fronto-central regions approximately 900 ms after stimulus onset. This effect was accompanied by a denser and more globally efficient directed connectivity network, suggesting increased large-scale coordination during the access of remote memories.

Although early prefrontal activation has been consistently associated with controlled search processes during AM retrieval [43, 44], prior works have also shown that spectrum power predicts successful retrieval in the searching period [45, 15]. In this study, we examined both early and late portions of the access period. Given that the minimum access duration was 1.9 s, we focused on the first 1500 ms after stimulus onset and the 1500 ms preceding the response to minimize contamination from motor preparation and elaboration processes. Notably, we identified reliable markers of memory remoteness only in stimulus-locked analyses. The absence of effects in response-locked data likely reflects the influence of motor and decision-related processes, which may temporally obscure earlier cognitive differences. These findings suggest that remoteness-related effects are specifically expressed during the early stages of memory access.

*θ*-band activity has been widely linked to hippocampal–cortical interactions and internally directed attention during memory retrieval [46]. The increased *θ* power observed for Remote AMs is consistent with prior findings showing modulation of *θ* activity during autobiographical retrieval [46, 47], including fronto-central effects around 1000 ms [18]. Importantly, *θ* power was not associated with subjective ratings of detail or importance, but instead showed a positive relationship with retrieval difficulty. This suggests that fronto-central *θ* activity reflects the cognitive demands of memory access rather than the phenomenological richness of the retrieved content.

Behavioral results further support this interpretation. Remote memories were rated as less detailed but more important than recent ones, consistent with the literature on memory transformation over time [19, 48, 6]. As autobiographical memories age, they tend to lose specificity and become more schematic, relying increasingly on semanticized representations [20, 49]. This process of semanticization may contribute why Remote memories may require greater cognitive effort during access, despite retaining their subjective importance. Importantly, our results do not allow us to determine whether these effects are driven by differences in memory content (episodic vs. semantic), but rather point to differences in the cognitive demands of memory access. Moreover, the absence of differences in access duration between conditions further suggests that response time may not provide a direct proxy for retrieval effort in this paradigm. In self-paced tasks, response times likely reflect a combination of processes, including decision thresholds and response strategies, which may obscure transient differences in cognitive demand. In contrast, neural measures such as *θ* activity may capture fine-grained variations in processing demands that are not reflected in overall response duration.

Our connectivity analysis provides further insight into these processes. During the access of Remote AMs, we observed a denser and more efficient *θ*-band network, suggesting enhanced large-scale integration. In contrast, Recent AMs were characterized by more locally clustered connectivity, consistent with the retrieval of vivid episodic details that may rely on more focal processing [22, 50]. Despite these differences, both conditions exhibited a predominant anterior-to-posterior flow of information, indicating a shared large-scale organizational pattern during AM access. This anterior-to-posterior flow aligns with previous findings implicating frontal regions in initiating memory search and posterior regions in reconstructing perceptual and contextual details [2, 3]. Moreover, frontal midline *θ* activity has been associated with cognitive control, monitoring, and self-referential processing, all of which are critical for autobiographical retrieval. Our findings extend this framework by showing that remote memories engage a more integrated version of this network, possibly reflecting a more effortful and generative mode of access [22].

These results are also consistent with theoretical accounts suggesting that Remote memories rely more heavily on distributed cortical representations and top-down processes [51]. In this context, increased *θ* synchronization may facilitate long-range communication between brain regions involved in imagery, emotional processing, and multimodal integration. Thus, *θ* activity may serve as a neural mechanism supporting the coordination of distributed processes required for accessing remote autobiographical memories.

An important consideration is that memory remoteness may be partially confounded with the quality of the mnemonic trace. Previous work has shown that vividness plays a critical role in shaping the neural dynamics of autobiographical retrieval. For instance, Sheldon and Levine (2013) reported that vivid memories are associated with an early and rapid recruitment of frontal, parietal, and limbic regions, consistent with a more direct retrieval process driven by imagery. In contrast, non-vivid memories exhibited a later and more spatially distributed pattern of activation, reflecting a more effortful and reconstructive retrieval process [22]. In this context, the increased *θ* power and more integrated network observed for Remote AMs in the present study may not be solely attributable to their temporal distance, but also to differences in trace quality, such as reduced vividness and increased reliance on generative processes. This interpretation is further supported by our behavioral findings, showing that remote memories were rated as less detailed and more difficult to access. Together, these results suggest that remoteness-related neural effects may, at least in part, reflect differences in mnemonic richness and retrieval demands rather than time per se.

Importantly, these findings depend on the temporal criteria used to define memory remoteness. By contrasting memories from the last year with those older than three years, we aimed to maximize differences in cortical reorganization. This approach is consistent with meta-analytic evidence showing that memory representations shift from hippocampal-dependent to cortical networks over time [23]. Our results suggest that such reorganization is detectable at the level of fast neural dynamics, emerging within the first second of memory access.

In summary, our results reveal qualitative differences in the neural mechanisms underlying access to recent and remote autobiographical memories. Remote memories are associated with increased *θ* power and more integrated network dynamics during early access, reflecting greater cognitive demands and large-scale coordination. By combining high-temporal-resolution EEG with directed connectivity analyses, this study provides new insight into the neural dynamics supporting autobiographical memory retrieval.

### 4.1 Limitations

Several limitations should be considered. First, scalp EEG does not provide direct access to deep brain structures such as the hippocampus, limiting our ability to assess hippocampal–cortical interactions directly. Second, the number of trials was relatively low and slightly unbalanced across conditions, which may affect the signal-to-noise ratio and the stability of connectivity estimates. Third, the self-paced nature of the task introduces temporal variability, although it enhances ecological validity. Finally, Granger Causality relies on weak stationarity assumptions that are only approximately satisfied in neural data; therefore, our results should be interpreted as time-resolved estimates of directed influence rather than definitive causal interactions. Future work combining EEG with source reconstruction or multimodal approaches could help address these limitations.

## 5 Funding information

This work was supported by a grant from the Agencia Nacional de Promoción Científica y Tecnológica PICT 2020 - 00956, the University of Buenos Aires UBACYT 2023 (20020220400164BA) and CONICET PIP 2021-2023.

## 6 Data availability

Data will be available after publication. o The datasets generated during and/or analysed during the current study are available from the corresponding author on reasonable request.

## 7 Author contributions

MCN: Conceptualization, Data collection, Formal analysis, Investigation, Methodology, Visualization, Software, Writing - original draft, Writing - review and editing. IF: Data collection, Resources. RSF: Conceptualization, Methodology, Resources, Funding acquisition, Writing - review and editing. MEP: Funding acquisition, Resources, Writing - review and editing. LB: Supervision, Investigation,Conceptualization, Methodology, Resources, Funding acquisition, Writing - review and editing.

## 8 Additional information

The authors declare no competing interests.

## Notes

### Competing Interest Statement

The authors have declared no competing interest.

### Summary of Updates

New analysis and results have been added.

